# Parallel evolution in the integration of a co-obligate aphid symbiosis

**DOI:** 10.1101/2020.01.14.905752

**Authors:** David Monnin, Raphaella Jackson, E. Toby Kiers, Marie Bunker, Jacintha Ellers, Lee M. Henry

## Abstract

Insects evolve dependencies - often extreme - on microbes for nutrition. These include cases where insects harbour multiple endosymbionts that function collectively as a metabolic unit. How do these metabolic co-dependencies originate, and is there a predictable sequence of events leading to the integration of new symbionts? While dependency on multiple nutrient-provisioning symbionts has evolved numerous times across sap-feeding insects, there is only one known case of metabolic co-dependency in aphids, between *Buchnera aphidicola* and *Serratia symbiotica* in the Lachninae subfamily. Here we identify three additional independent transitions to the same co-obligate symbiosis in different aphids. A comparison of recent and ancient associations allows us to investigate intermediate stages of metabolic and physical integration between the typically facultative symbiont, *Serratia*, and the ancient obligate symbiont *Buchnera*. We find that these uniquely replicated evolutionary events support the idea that co-obligate associations initiate in a predictable manner, through parallel evolutionary processes. Specifically, we show (i) how the repeated losses of the riboflavin pathway in *Buchnera* leads to dependency on *Serratia*, (ii) evidence of a stepwise process of symbiont integration, whereby dependency evolves first, then essential amino acid pathways are lost (at ~30-60MYA), which coincides with increased physical integration of the companion symbiont; and (iii) dependency can evolve prior to specialised structures (e.g. bacteriocytes), and in one case with no direct nutritional basis. More generally, our results suggest the energetic costs of synthesising nutrients may provide a unified explanation for the sequence of gene loses that occur during the evolution of co-obligate symbiosis.

## RESULTS

### Independent transitions to co-obligate dependency

Sap-feeding insects have provided elegant case studies of the evolution of co-obligate dependencies, whereby organisms harbour multiple endosymbionts that function collectively as a metabolic unit. These include species of mealybugs that depend on endosymbionts, which in turn harbour their own endosymbionts, and cicadas, in which one symbiont has fragmented into distinct but interdependent lineages [1–5]. What processes drive multiple microbial species to join into metabolic co-dependencies [6], and more generally, is there a predictable, deterministic sequence of events leading to the genomic and physical integration of new symbionts?

The aphids are an ideal lineage to study early stage co-obligate dependencies. The majority of aphid species harbour a single obligate symbiont, *Buchnera aphidicola*, and a second non-obligate symbiont *Serratia symbiotica* (hereafter referred to as *Buchnera* and *Serratia* for simplicity). While *Serratia* is found at intermediate frequencies in numerous aphid species, the symbiont has transitioned to a co-obligate relationship with *Buchnera* in the Lachninae subfamily [7–9]. Such co-obligate functioning is marked by *Buchnera* losing metabolic capabilities, namely the ability to synthesise the essential nutrients riboflavin and secondarily tryptophan [10]. Our aim was to determine if: (i) other cases of obligate co-dependencies have arisen across the aphids, and (ii) to ask whether these transitions followed predictable genomic, metabolic and physical trajectories. Such patterns can provide insight into the evolutionary processes that have led to the genome structure of more ancient multi-partner symbioses [6].

Using data on the symbiont prevalence in 131 aphid species from [11], we identified species that carry *Serratia* at a high frequency, and then tested aphid populations in both the United Kingdom and the Netherlands for obligate dependency on the symbiont. We defined species as having evolved obligate reliance on *Serratia* if, i) all individuals within populations carry the symbiont, and ii) they experience a significant fitness reduction when the symbiont is removed. We screened for the presence of *Serratia* using PCR and measured dependency by “curing” individual aphids with antibiotics that selectively removed *Serratia* without affecting *Buchnera*, and then determining the lifetime fecundity of the aphids in the presence and absence of the symbiont.

We identified ubiquitous *Serratia* symbioses in seven aphid species (Table S1), representing three independent co-obligate transitions in *Microlophium carnosum, Aphis urticata*, and in the *Periphyllus* genus. In the *Periphyllus* genus, *Serratia* was consistently present in five species we surveyed, and we confirmed obligate dependency via curing *Serratia* in both *P. hirticornis* and *P. lyropictus*. This suggests a single transition into an obligate relationship with *Serratia* at the origins of the *Periphyllus* genus (see below). Curing had the most dramatic effect in species of the *Periphyllus* genus, potentially reflecting a longer-term evolutionary association with *Serratia* (Fig. 1). We confirmed the antibiotic treatments had no significant effect on the fecundity of our control aphid species *Acyrthosiphon pisum*, which harbours *Serratia* as a facultative symbiont and the uninfected *Macrosiphoniella artemisiae* (Fig. 1A, Table S2, Fig. S1, Table S3). Likewise, we confirmed with quantitative PCR that the antibiotic treatment reduced *Serratia* density (Fig. 1B, Table S4), but did not reduce *Buchnera* density (Fig. 1C, Table S5).

**Figure 1.**
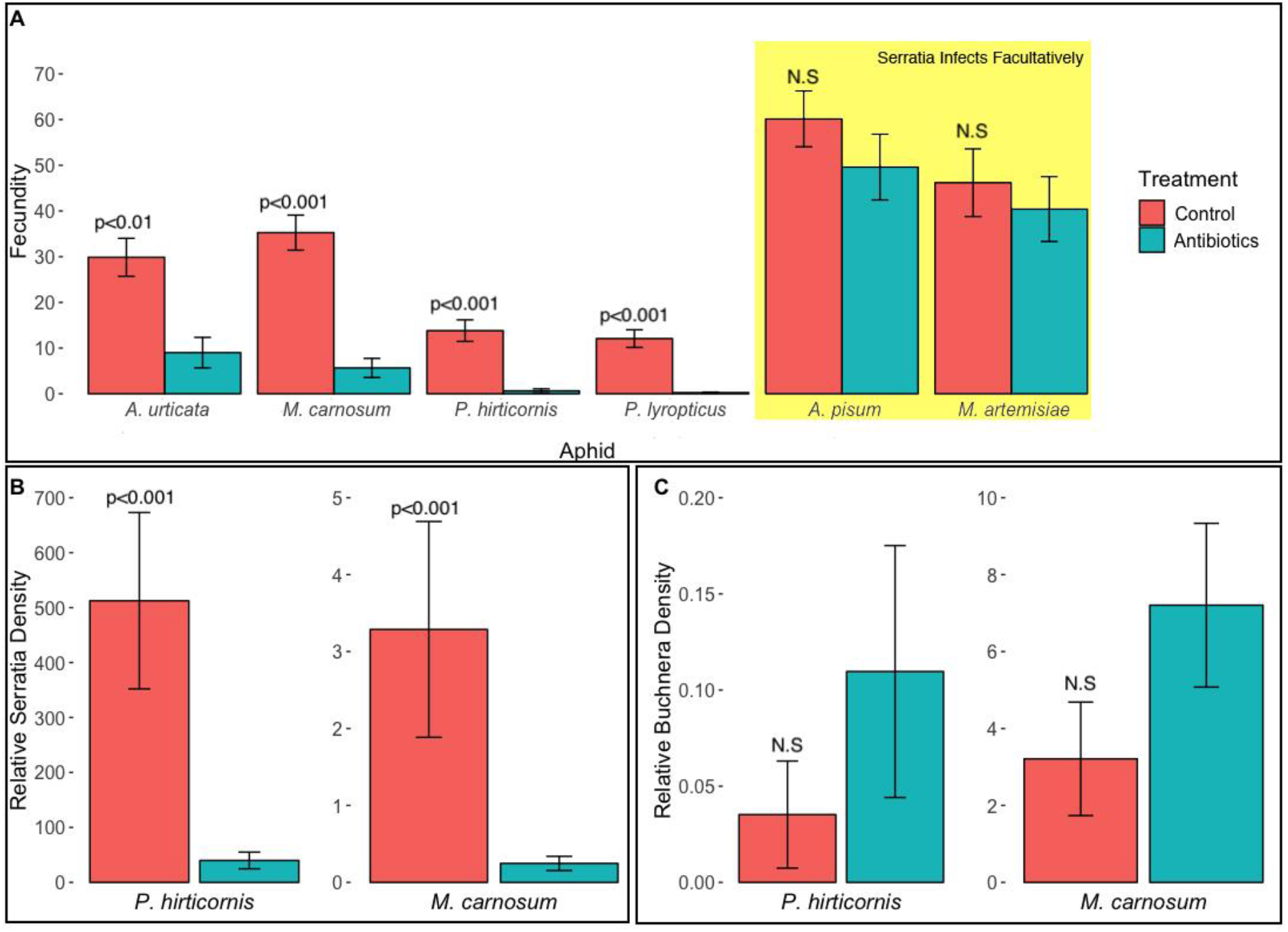
Antibiotic curing of *Serratia* in aphids. Effect of antibiotic curing on aphid lifetime fecundity (A) in aphid species representing three independent evolutionary transitions to co-obligate dependency (*A. urticata*, *M. carnosum*, and *P. hirticornis* and *P. lyropictus* representing the *Periphyllus* genus). The yellow box highlights species that host *Serratia* as a facultative symbiont. Results synthesised from two independent trials on aphids from the United Kingdom and the Netherlands. Effect of antibiotic curing on *Serratia* density (B) and *Buchnera* density (C) compared to the density of host cells.

We next estimated the origins of obligate dependency on *Serratia* using deep coverage 16S amplicon sequencing from our field collected populations, and previous data on *Serratia* associations in aphids [11]. First, we found evidence of a more ancient relationship between *Serratia* and aphids in the *Periphyllus* genus: amplicon sequencing confirmed *Serratia* was absent from *Chaitophorus* aphids, a sister lineage to the *Periphyllus* clade (Fig. S2). This suggests co-dependency originated at the divergence of these two genera, an estimated 63-79MYA ago (Fig. 2, Fig. S3). Second, we found evidence of more recent origins of *Serratia* obligate dependency (<30MYA) in *A. urticata* and *M. carnosum*. Specifically, *Serratia* was either absent, or present only as a facultative infection, i.e. only present in some individuals, in the related species of *A. idaei* and *A. fabae* (for *A. urticata*). Lack of obligate dependency was likewise confirmed in *M. euphorbiae*, and *A. pisum*, related to *M. carnosum*.

**Figure 2.**
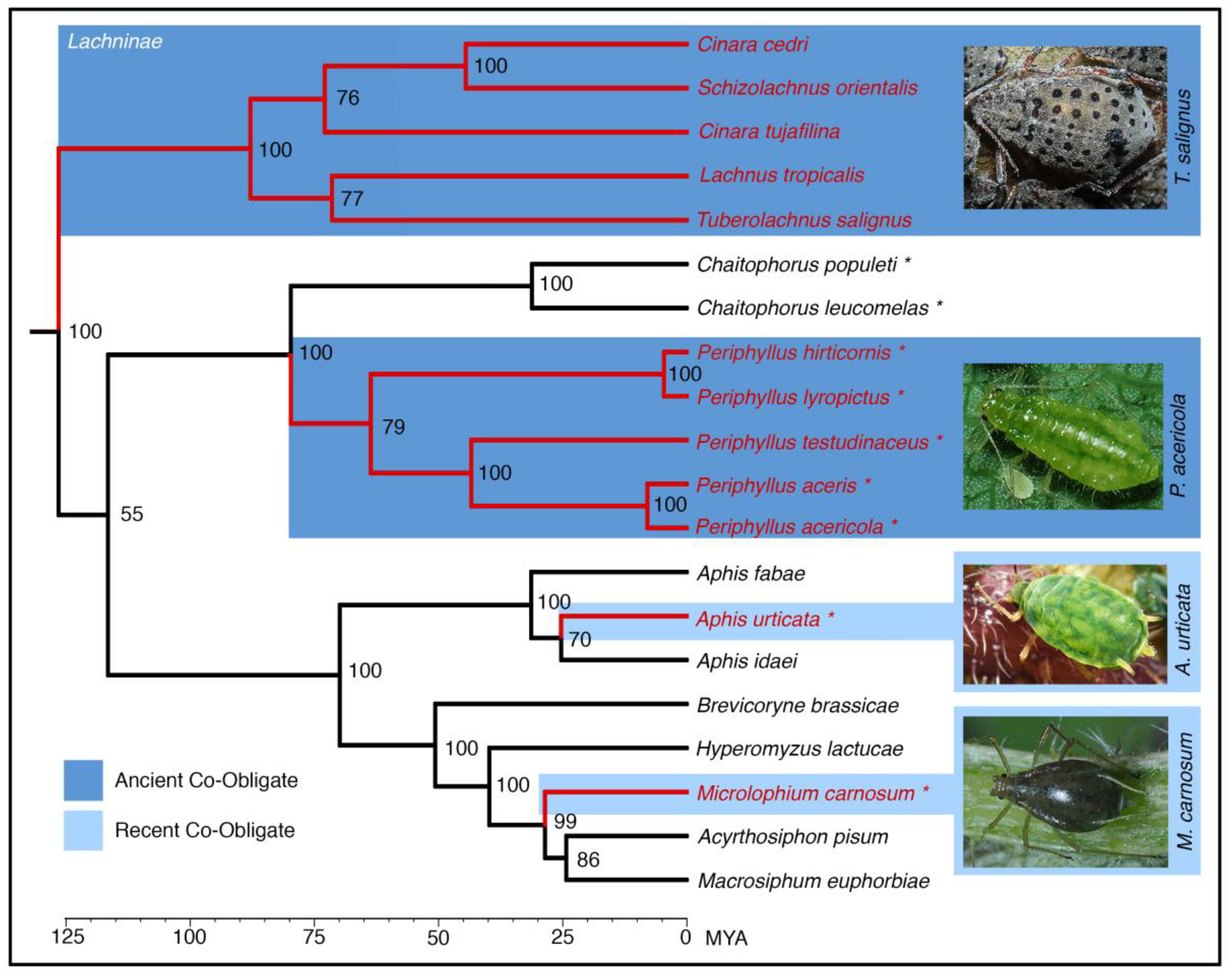
A dated aphid phylogeny showing lineages with co-obligate symbioses. Branch colours represent aphid lineages that have evolved obligate dependency on *Serratia* (red) or those that carry the symbiont facultatively (black). Light blue shading represents more recent co-obligate associations, and dark blue represents ancient associations for the entire clade of host species. Asterisks denote data from this study. Other data is compiled from several studies [11–13]. Numbers on the nodes of the tree are bootstrap values. Node ages (MYA) and confidence intervals in Fig S3. Photos from InfluentialPoints.com.

### Genomic basis of *Buchnera-Serratia* metabolic complementarity

We then asked if evolving obligate dependency on *Serratia* was associated with a consistent genomic signature in aphids, and more specifically whether *Buchnera-Serratia* metabolic complementarity originates in a predictable manner across host lineages. We obtained whole genome sequencing data for *M. carnosum, A. urticata* and three *Periphyllus* species. We then used previous published data from the Lachninae subfamily (*Cinara cedri*, *Cinara tujafilina* and *Tuberolachnus salignus*) to compare the gene losses in *Buchnera* from the four independent transitions into an obligate relationship with *Serratia*. This included the three new cases of co-obligate dependency identified here, and the previously identified cases in the Lachninae subfamily. Our analysis centred on pathways and genes involved in essential nutrients provisioning to the host (Table S6). Specifically, we focused on pathways which have experimental evidence for being essential for the aphid: riboflavin [14] and essential amino acids [15–21]. Of particular interest was the riboflavin pathway in *Buchnera*, as the loss of this pathway has been hypothesized to trigger the dependency on *Serratia* in the Lachninae aphids [10].

We found a consistent signature for the loss of *Buchnera*’s riboflavin pathway in both *M. carnosum* and aphids in the *Periphyllus* genus (Fig. 3). In *M. carnosum, Buchnera* is missing one gene, part of the ribD complex, which is essential to the riboflavin pathway. In the *Periphyllus* genus, by contrast, the full pathway is missing, as it is in the Lachninae subfamily. Previous work in the Lachninae aphids suggests that *Buchnera* has also secondarily lost the capacity to synthesise the amino acid tryptophan in certain species (e.g. *C. cedri* and *T. salignus*). We find similar losses in the *Periphyllus* lineage, where the majority of genes in the tryptophan pathway have either been lost or pseudogenised and complementary gene copies have been retained in the *Serratia* genome (Fig. 3). Conversely, the tryptophan pathway has been retained in the *Buchnera* genomes of both *M. carnosum* and *A. urticata*, the more recent co-obligate relationships. This result suggests an advanced stage of functional losses in *Buchnera* of *Periphyllus* aphids, further supported by losses in several additional amino acid pathways that now also appear to have been taken over by *Serratia*.

**Figure 3.**
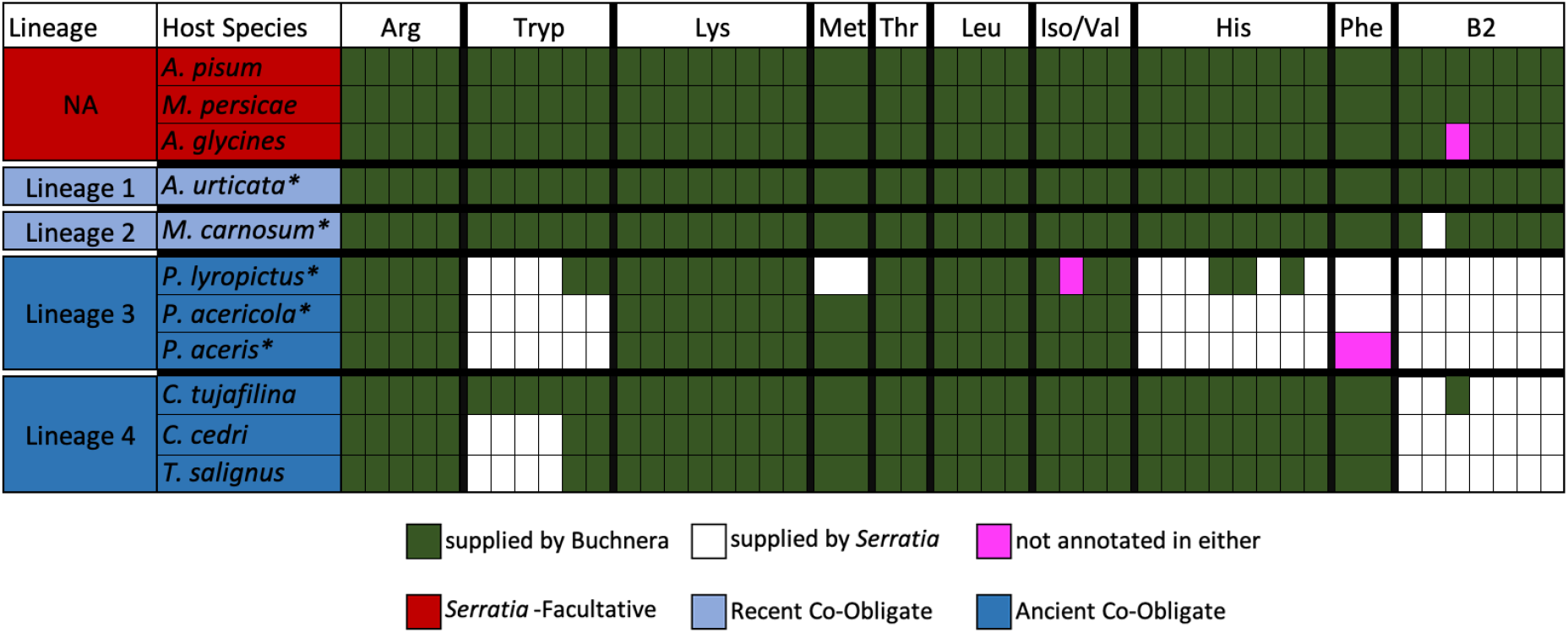
Essential nutrient-provisioning genes of *Buchnera* and *Serratia* genomes. Aphid lineages that have evolved dependency on *Serratia* are numbered. Coding capabilities of symbionts for essential amino acids and riboflavin are presented for each aphid species. Each cell represents a single gene with colour coding depicting whether the gene is present in the genome of *Buchnera* (green) or *Serratia* (white). Gene content of *Buchnera* from *A. pisum*, *M. persicae* and *A. glycines* are included as a comparison to species where *Buchnera* is the sole obligate symbiont. Gene names and additional information, including gene redundancies, can be found in Table S6. Asterisks denote genomic data from this study.

In contrast, *Buchnera* has retained the complete pathways to synthesise all of these essential nutrients in *A. urticata*. This is surprising given the consistency of gene losses in *Buchnera* of *M. carnosum*, aphids in the *Periphyllus* genus, and the Lachninae aphids – all which are co-obligately dependent on *Serratia. (Acyrthosiphon pisum, Myzus persicae* and *Aphis glycines*) the co-obligate association of *Serratia* and *Buchnera* in *A. urticata* has only six genes missing in *Buchnera*, in which there are complementary gene copies in *Serratia* (see Table S7 for more detail). None of these six genes have direct links to essential nutrient pathways (see Table S7 for more detail). This suggests co-obligate dependency can arise in this system through alternative starting points, including non-nutritional pathways.

General genomic features of the different *Buchnera* strains likewise support the hypothesis that the co-obligate *Serratia* symbioses found in Lachninae and *Periphyllus* aphids are more ancient compared to *M. carnosum* and *A. urticata* (Fig. S4). Both the genome size and GC content of *Buchnera* are highly reduced in the Lachninae and *Periphyllus* clades, suggestive of a more advanced degree of degradation and integration with *Serratia*. Gene redundancies are also indicative of the age of the co-obligate associations. In *M. carnosum* and *A. urticata*, the genomes of *Serratia* and *Buchnera* still both contain a significant number of the same genes involved in synthesising nutrients that are essential for the host aphid (72.5% and 39.2% respectively). Conversely, in the *Periphyllus* lineages, both *P. acericola* and *P. aceris* have only 11.5% gene redundancy between the two symbionts, whereas in *P. lyropictus* there is 47.1%. The latter increase is likely due to *Serratia* being recently replaced by another *Serratia* strain within this aphid lineage.

### Physiological integration of *Serratia* in co-obligate symbiosis

Lastly, we studied the physical integration of symbionts with their host - and each other - to look for physical signatures of co-obligate symbiosis. We expected that a greater degree of metabolic reliance on *Serratia* in the *Periphyllus* aphids would correspond with greater physiological integration, for example through an increase in specialised cells (bacteriocytes) to house *Serratia*, and its abundance within the aphid is expected to increase.

We performed Fluorescent *in situ* Hybridisation using probes specifically targeting *Buchnera* and *Serratia*. As predicted, we found increased physiological integration of *Serratia* in host lineages corresponding with a greater reliance on *Serratia*. Higher integration is likely driven by the need to compensate for metabolic losses in *Buchnera*. In the most extreme case, we found that the *Periphyllus* aphids evolved a large organ (bacteriome) containing numerous bacteriocytes to house *Serratia* in their abdomen (Fig. 4). In line with previous predictions, we also found that the ratio of *Serratia* to host genome copies increased dramatically to an abundance ratio of 512:1 in *P. hirticornis*. This is compared to a 3:1 ratio found in the less integrated co-obligate of *M. carnosum* (Fig. 1B).

**Figure 4.**
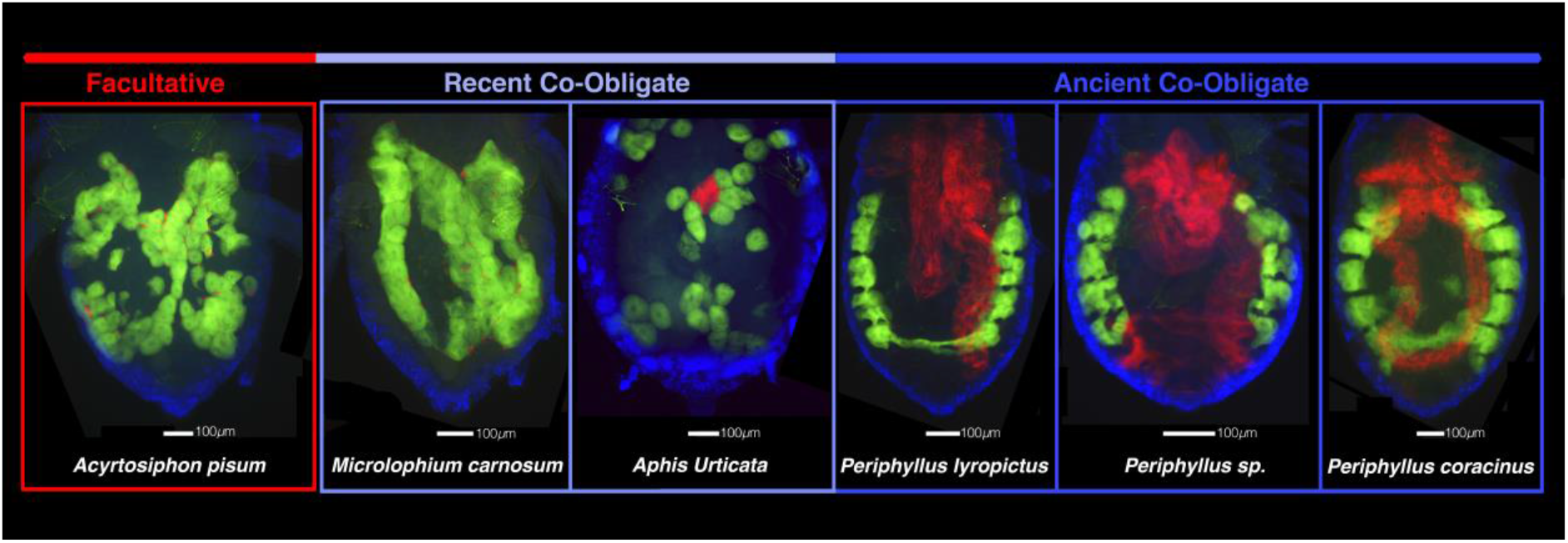
FISH images of abdomens from 6 aphid species. *Buchnera* and *Serratia* are highlighted in green and red, respectively. The coloured bar represents the degree of reliance on *Serratia* - facultative (red) or co-obligate (blue). The shades of blue represent the degree of physiological integration with the host, from the least (*M. carnosum*) to the greatest (*Periphyllus* spp.).

In contrast to high physical integration in the *Periphyllus* aphids, we found that both *M. carnosum* and *A. urticata* exhibit minimal physiological integration of *Serratia*. In *A. urticata, Serratia* is localised in a small cluster of relatively large cells (~4) forming a small bacteriome surrounded by *Buchnera*-containing bacteriocytes. In *M. carnosum*, *Serratia* is the least integrated, with the symbiont being localised in sheath cells surrounding the *Buchnera*-containing bacteriocytes. This pattern is similar to the one found in *A. pisum* where *Serratia* maintains a consistently facultative relationship with its host.

## DISCUSSION

Dependency on multiple co-dependent symbiotic microbes has originated numerous times in the evolution of eukaryotes. However, the vast majority of co-obligate symbioses are ancient. Data on recent associations are needed to reveal the evolutionary processes that initiate co-dependency, and provide insight into the intermediate steps leading to the extreme genomic and physiological integration observed in ancient associations. By comparing ancient and recent associations in aphids, we find strong evidence that the mechanisms initially binding symbiotic partners in obligate relationships occur in a deterministic, predictable manner. Specifically, we find that dependency on *Serratia* originates through parallel evolutionary trajectories marked by repeated losses of the same nutrient pathways in *Buchnera* across multiple host lineages. Our genomic and physical data shows stepwise processes of symbiont integration, with the losses of essential amino acid pathways occurring between 30-60MY after co-dependency evolves. This is followed by a second phase of dependency characterised by greater physical integration of *Serratia* in the more ancient obligate partnerships of the *Periphyllus* genus compared to more recently adopted co-obligate associations of *A. urticata* and *M. carnosum*.

Our results provide the first evidence that *Buchnera* has repeatedly lost the capacity to produce the essential nutrient riboflavin in multiple aphid lineages. In each case, where the pathway to synthesise riboflavin has been lost, *Serratia* has retained genes to compensate for these metabolic changes in *Buchnera*. In several species within the Lachninae sub-family, the tryptophan pathway is also missing, suggesting that once the co-obligate symbiosis with *Serratia* is established this amino acid is secondarily lost in *Buchnera* [10]. Our work confirms that the tryptophan pathway has likewise been lost in *Periphyllus* aphids, and that the capacity to synthesise this amino acid is vulnerable to deletion. Tryptophan is one of the most costly essential amino acid to synthetise [22], and it has been hypothesised that this may explain why its lost is associated with the presence of a second obligate symbiont [6]. The second most energetically expensive amino acid synthesis pathways (phenylalanine, histidine, methionine and isoleucine/valine) are likewise lost in the *Buchnera* of *Periphyllus* aphid. This is in line with past work documenting the loss of energetically expensive amino acids, and complementation by a companion symbiont, in several ancient co-obligate symbioses, including *Sulcia* of some Auchenorrhyncha families (e.g. spittlebugs and cicadas) and in *Carsonella* the primary symbiont of Psyllids. In *Sulcia*, the amino acid pathways appear to have been lost around 60MY after co-dependency evolved, prior to the common ancestor of cicadas, sharpshooters and spittlebugs [23]. We find amino acid pathways are only lost in *Buchnera* in the more ancient co-obligate associations of the *Periphyllus* and in some Lachninae species. In contrast, *Buchnera* has retained these functions in the more recent co-obligates of *A. urticata* or *M. carnosum*. This suggests essential amino acids are only susceptible to deletions in the second phases of losses once selection has been relaxed by the presence of a new obligate symbiont, approximately 30-60MY after co-dependency evolves. These results also provide strong support for the hypothesis that the energetic costs of synthesising nutrients may provide a unified explanation for the sequence of gene losses that occur during the evolution of co-obligate symbiosis (Table S8).

In the case of *A. urticata*, we find that dependency on *Serratia* has evolved with no direct link to nutrient provisioning. This may be a case of “evolved dependency” where an organism becomes so adapted to the constant presence of a partner that is loses the ability to perform well on its own [24,25]. This scenario has been proposed to explain the dependency on *Wolbachia* in the Hymenoptera *Asobara tabida* [26,27] and the fitness-enhancing effect of grazing on plants [25]. This suggests that tripartite obligate symbioses, at least in some cases, may arise through processes other than degradation of the ancestral symbiont - as typically thought - but rather dependency may evolve first and then deterioration follows.

More generally, our data support the hypothesis that lineages which recently acquired co-obligate symbionts will have cases of overlapping gene complexes. This is seen in both *A. urticata* and *M. carnosum*, where *Serratia* shares many redundant genes with *Buchnera* in pathways for essential nutrients synthesis. While two of the three *Periphyllus* species showed high degrees of metabolic complementation, there was a single case (*Periphyllus lyropictus*) in which we did not document this pattern. This is likely due to *Serratia* being replaced within this lineage. Symbiont replacement is an important mechanism by which maladaptive symbionts are replaced with new functional ones [28,29]. In the Lachninae aphids, *Serratia* has been replaced on multiple occasions, including by other more recently acquired *Serratia* strains [9,12,13].

The multiple independent origins of *Buchnera-Serratia* co-obligate symbioses also provide a unique opportunity to study the evolution of physical integration of host and co-obligate symbionts. Theory suggests that more ancient associations should be characterised by greater integration [30], both in terms of housing structures and symbiont densities within hosts. Such integration is expected as the new symbiont takes a more metabolically demanding role. While our results generally support this hypothesis, we found that the evolution of specialised structures to house *Serratia* differed between the two most ancient aphid lineages. In the Lachninae, both co-obligate symbionts are thought to be located within the same bacteriome, although the arrangement of bacteriocytes differ among species [12]. The *Periphyllus* aphids, by contrast, house the two symbionts in separate structures. These configurations potentially represent alternative solutions to the same problem: allowing *Serratia* to reach densities high enough to efficiently perform its nutrient-provisioning role. Our curing data further support the idea that greater integration coincides with more reliance on *Serratia* in ancient co-obligate, as symbiont removal had the most dramatic fecundity effect in the *Periphyllus* genus.

In the cases of *A. urticata* and *M. carnosum*, physical and genomic integration data suggest *Serratia* is a more recently adopted co-obligate association. In *A. urticata*, *Serratia* is housed in a single small bacteriome. In *M. carnosum*, *Serratia* is not hosted in primary bacteriocytes at all, but rather in sheath cells that surround the large *Buchnera* containing cells – a potential indication of its relatively recent transition to an obligate symbiont. According to this hypothesis, this lack of *Serratia* integration is indicative of its recent role as a facultative symbiont. These findings also suggest that - at least in some cases - dependency evolves prior to the evolution of specialised structures to house symbionts.

Studying evolutionary transitions to obligate and co-obligate symbiosis is difficult because most events are characterized by single and ancient origin across large, diversified clades. This makes comparisons with outgroups less informative and prevents testing ideas on the relative importance of deterministic versus stochastic processes. The uniquely replicated evolutionary events of *Buchnera-Serratia* co-obligate symbiosis in different aphid lineages provides a degree of temporal resolution that demonstrates co-obligate associations can form in a predictable manner. Furthermore, our findings indicate that genomic integration may occur prior to physiological integration. Our results provide evidence that the evolutionary forces that bind multiple organisms into a single metabolic unit operate by deterministic stepwise processes, which allows us to better understand the role of symbioses in the evolution of complex organisms.

## Supporting information

Supplemental information, Main

## ACKNOWLEDGEMENTS

The authors thank Alejandro Manzano-Marín for consultation on optimal parameters for genome assembly, Mario Dos Reis Barros for advice on molecular clocking. This project was funded by LH NERC IRF (NE/M018016/1), Marie Curie (H2020-MSCA-IF-2017-796778-SYMOBLIGA), JE NWO VICI (865.12.003) and ETK (Ammodo Funds).

## AUTHOR CONTRIBUTIONS

Conceptualisation, D.M., L.H., and R.J.; Methodology, D.M., L.H., M.B., and R.J.; Software, R.J.; Validation, D.M., L.H., M.B., and R.J.; Formal Analysis, D.M., M.B., and R.J.; Investigation, D.M., L.H., M.B., and R.J.; Resources, D.M., L.H.; Data Curation, R.J.; Writing - Original Draft, D.M., L.H., R.J., and T.K.; Writing - Review & Editing, D.M., J.E., L.H., M.B., R.J., and T.K.; Visualisation, R.J.; Supervision, J.E. and L.H.; Project Administration, L.H.; Funding Acquisition, D.M., L.H.

## STAR METHODS

## LEAD CONTACT AND MATERIAL AVAILABILITY

This study did not generate new unique reagents. Further information and requests for resources should be directed to the Lead Contact, Lee Henry.

## EXPERIMENTAL MODEL AND SUBJECT DETAILS

The following aphid species were used in this study: *Acyrthosiphon pisum*, *Macrosiphoniella artemisiae*, *Aphis urticata*, *Microlophium carnosum*, *Chaitophorus leucomelas*, *C. populeti*, *Periphyllus acericola, P. aceris, P. coracinus*, *P. hirticornis*, *P. lyropictus* and an unknown *Periphyllus* species (referred to as *Periphyllus sp*. in the main text). Clonal lines of aphids were maintained in the lab at 15°C with a 16 hours light 8h dark UV light cycle on a leaf of their host plant: *Vicia faba* (*A. piusm*), *Artemisia vulgaris* (*M. artemisiae*), *Urtica dioica* (*A. urticata* and *M. carnosum*), *Acer spp*. (*Periphyllus spp*.) embedded in agar in a Petri dish to keep the leaves fresh. Leaves were changed as needed.

## METHOD DETAILS

### Assessment of *Serratia* prevalence

#### Collection and identification of aphid samples

Aphids were collected in the UK and the Netherlands between 2011 and 2019. They were dislodged by beating plants over a white tray or removed manually from the plant, before being placed in 100% ethanol or collected live for curing experiments. Resampling of the same aphid clones was minimised by separating collections from the same plant species by at least 10 m. Aphids were identified by barcoding based on data from [31] and morphological examination following (Heie and others 1980-1995). We sequenced the COI barcoding region using standard protocols for DNA extraction, amplification and editing and performed alignments with MUSCLE in CODONCODE ALIGNER version 4.0.2 (CodonCode Corporation 2012, Centerville, MA, USA). Genomic DNA was extracted from individual specimens using DNeasy Blood and Tissue kits (Qiagen, Venlo, Netherlands) and we amplified an approximately 700 bp DNA fragment of the cytochrome c oxidase I (COI) mitochondrial gene using Lep F (5’-ATTCAACCAATCATAAAGATATTGG-3’) and Lep R (5’-TAAACTTCTGGATGTCCAAAAAATCA-3’) primers, which was sequenced in the forward direction. Aphids were identified to species by comparing COI sequence data to the online databases BOLD (http://www.barcodinglife.org) and GenBank using BLAST. Morphological examination was carried out by macerating individual aphids in KOH (Potassium Hydroxide) and mounting them on microscope slides.

#### Diagnostic screening for *Serratia symbiotica*

We confirm all *A. urticata, M. carnosum* and five species of *Periphyllus* aphids ubiquitously carry *Serratia*, in the UK and the Netherlands, by amplifying a partial region of the 16S rRNA gene using the specific primers 16sS A1 (5’-AGAGTTTGATCMTGGCTCAG-3’) and 16sS 2R (5’-TTTGAGTTCCCGACTTTAACG-3’) and sequencing the amplicons. The PCR cycling conditions were as follow: 3 minutes at 95°C followed by 30 cycles of 30 seconds at 95°C, 1 minute at 52°C and 1 minute at 72°C, and finally 5 minutes at 72°C. To confirm the primers were only amplifying *S. symbiotica* we compared sequences to published records on GenBank using BLAST. If the specific primers failed to amplify due to primer binding specificity we used more general *Serratia* primers that amplify diverse *Serratia* species: 16sS 10F (5’-AGTTTGATCATGGCTCAGATTG-3’) and 16sS R443R (5’-CTTCTGCGAGTAACGTCAATG-3’), and confirmed the presence of *S. symbiotica* by compared sequences to those on GenBank using BLAST. PCR cycling conditions were as follows, 2 minutes at 94°C, followed by 10 cycles of one minute at 94°C, 1 minute at 65°C-55°C (touchdown in 1°C steps) and 2 minutes at 72°C 2:00, followed by 25 cycles of 1 minute at 94°C, 1 minute at 55°C and 2 minutes at 72°C, and finally, 6 minutes at 72°C [33].

To estimate the origins of the co-dependency in *Periphyllus* aphids we sampled for *Serratia* in the *Chaitophorus* genus, which is a sister taxon to *Periphyllus* [11]. We deep-sequenced the universal bacterial 16S rRNA gene in *Chaitophorus populeti* (7 samples) and *C. leucomelas* (1 sample) to confirm *Serratia* was not ubiquitously present in these aphids, indicating they had not evolved co-dependency on the symbiont or been replaced. We PCR amplified the V4 region of the bacteria 16S rRNA gene following standard protocols [34], and deep-sequenced the amplicons using the Illumina MiSeq2000 platform. 16S rRNA analysis was preformed using the standard operating procedure for MOTHUR [35], including read joining filtering, and Operational taxonomic unit (OTU) selection at 99 percent. Taxonomic assignments of the reads were preformed using the full length SILVA alignment [36] available from mothur.org. As a final step samples were filtered using R to only consider OTUs at a one percent relative abundance or higher in the sample. No OTU(s) corresponding to *Serratia symbiotica* were found in any of these samples. The total and absolute number of OTU reads for each symbiont species are presented in Table S9.

### Curing experiments

To selectively cure aphids of *Serratia* while not affecting *Buchnera*, we used antibiotics that specifically target cell walls, which are absent in *Buchnera* [37,38]. Curing experiments were conducted on *A. urticata, M. carnosum*, and the two *Periphyllus* species that we were able to successfully culture in the lab: *P. hirticornis* and *P. lyropictus* in both the UK and the Netherlands. We were unable to culture the remaining three *Periphyllus* species in the lab so they were not include in the curing assays. *Acyrthosiphum pisum* (UK and NL) and *M. artemisiae* (UK only) were included to confirm the antibiotic treatments had no consistent negative fitness effects on species that harbours *Serratia* as a facultative symbiosis (*A. pisum*) or were uninfected by the symbiont (*M. artemisiae*). The antibiotic solution was obtained by mixing 10 mg/mL of Ampicillin sodium salt, 5 mg/mL Cefotaxime sodium salt, and 5 mg/mL Gentamicin in water. All antibiotics were obtained from Sigma-Aldrich (St. Louis, MO, USA). A single leaf of the host plant was cut and placed in a 0.5 mL Eppendorf tube filled with either one of two treatments i) the antibiotic solution or ii) water (control treatment). For the curing assay we placed 3-5 one or two day old aphid nymphs on a leaf and left them to feed for five (UK experiment) or three (Dutch experiment) days on either water or antibiotic solution. At the end of the treatment, aphids were transferred to their own individual Petri dishes, each containing a single leaf in agar from their appropriate host plants. The leaves were changed each week and the lifetime fecundity of each aphid was recorded. In the UK experiment, a sub-sample of first-generation aphids (9 control and 11 treated *M. carnosum* and 10 control and 10 treated *P. hirticornis*) were sacrificed at ~20 days old to quantify symbiont density using qPCR. The experiment was performed in the temperature and lighting conditions described previously.

### Quantitative PCR

The relative density of *Buchnera* compared to the density of *Serratia* was measured by quantitative PCR using three genes present as single copies in the aphid and each of the symbionts genomes from whole aphid DNA extractions (Table S10). The copy number of the symbiont genes in the DNA samples was measured by quantitative PCR on a CFX Connect Real-Time PCR Detection System (BioRad, Hercules, California, U.S.A.). The PCR reaction mixture included 10 μl Luna Universal qPCR Master Mix (New England BioLabs, Ipswich, Massachusetts, USA), 7 μl H_2_O, 0.5 μl of each primer (10 nM), and 2 μl DNA. The cycling conditions were: 15 min activation at 95°C followed by 40 cycles at 95°C for 15 s, at 60°C for 1 min, and 95°C for 15 s. The mean efficiencies were calculated using a ten-fold series of dilutions from 10^2^ to 10^7^ copies of purified PCR products. The efficiencies were 95.7 for the aphid gene, 96.8 for the *Buchnera* gene, and 97.3 for the *Serratia* gene. Duplicate samples were used for the determination of DNA quantities. As the deviations between the duplicates were below 0.5 cycles, the mean Cp values were used to calculate starting quantity. For each sample, the starting quantity for the *Buchnera* gene was divided by the starting quantity for the aphid gene to obtain the *Buchnera* density. Relevantly similar calculations were used to obtain the *Serratia* density.

### Phylogeny

The aphid phylogeny was build using a concatenation of four genes: *Elongation factor 1 alpha, 12S ribosomal RNA, 16S ribosomal RNA* and *Cytochrome oxidase subunit 1*. Sequences were obtained from NCBI (Table S11) and from the genomic data original to this study. Genes were concatenated using CLC genomic workbench 12.0 and aligned using MUSCLE [39]. Maximum Likelihood (ML) phylogeny was generated using the online PhyML server [40]. The phylogeny was bootstrapped 100 times, and rooted using sequences from *Adelges japonicus*, *A. couleyi*, *Candidatus* Ishikawaella capsulata and *Salmonella enterica*. The Adelgidae are basal to the Aphididae [41], *Candidatus* Ishikawaella capsulata and *Salmonella enterica* are outgroups belonging to the same family as *Buchnera aphidicola*. The tree was visualised using FigTree v1.4.4 [42]. *Chaitophorus saliniger* was included to improve node support then pruned from the phylogeny to only retain species for which *Serratia*-infection data are available.

We dated the phylogeny using the mcmctree function in PAML [43]. The calibrated the molecular clock using the estimated divergence (97.45-77.65 MYA) between the Lachnini (e.g. *T. salignus*) and Eulachnini (e.g. *C. cedri*) [44].

### Whole genomes sequencing, assembly and analysis

The genomes of individual aphids including their symbionts were sequenced at Centre for Genomic Research (University of Liverpool) from Qiagen DNeasy kit DNA extractions. The libraries were prepared using the Nextera XT kit, and sequenced on an Illumina HiSeq 4000 (paired-end, 2×150 bp reads). Seven samples were multiplexed on one lane. Average genome coverage for the endosymbionts of interest was ~950x for *Buchnera* and ~97x for *Serratia* (Table S12).

Reads were trimmed for quality and Illumina adaptors were removed using Trimmomatic [45] under default settings. Reads were assembled using SPAdes v3.11.1[46] in two stages. In the first stage and assembly was built, using assembly only mode with other parameters as default. The reads were then mapped back to this assembly using bwa mem. Contigs were partitioned into *Buchnera* and *Serratia* bins based on a DIAMOND [47] search of the contigs against the NCBI’s non-redundant Refseq protein database [48]. The reads mapping to the contigs in the *Buchnera* bin were then reassembled using SPAdes, this time using error correction and in careful mode with kmer sizes of 33, 55, 77, 99 and 127. The contigs of the resulting assembly were filtered by coverage and identity, then blasted against the NCBI’s non-redundant Refseq nucleotide database [48]. The results were manually inspected and contigs belonging to species other than *Buchnera* were removed.

Three of the five genomes, the *Buchnera* strains belonging to the *Periphyllus* species, were not able to be fully closed due to repetitive high AT content regions. These genomes were aligned against *Buchnera aphidicola* strain APS [49] and each other using MAUVE [50] and inspected using Geneious Prime 2019.06.17 (https://www.geneious.com) to ensure the contigs were oriented correctly. Using the high level of gene synteny between *Buchnera* lineages we manual inspected the gaps of all five genomes to confirm they could not contain genes relevant to provisioning of essential nutrients. In only one case, the genome of *P. aceris* contained a gap with the genes ribE and ribD, which are part of the riboflavin pathway. In this case, the remaining genes in the riboflavin pathway were confirmed to be absent followed by a comparison with the *Buchnera* of the other two *Periphyllus* species. Both species had a similar deletion that spanned the area of the gap. Finally, the assembled sequencing data from before the binning stage was inspected to ensure these genes were not overlooked due to incorrect binning. In this way we confirmed the pathway was non-functioning in this *Buchnera* lineage

We annotated both *Serratia* and *Buchnera* genes using DFAST [51] with Escherichia coli K-12 genome annotations as a reference [52]. To ensure we did not miss any genes in our final assemblies, we also annotated the initial assembly prior to partitioned into *Buchnera* and *Serratia* bins, and confirmed they did not contain any additional genes involved in nutrient provisioning. Genes making up each pathway were determined using the Metacyc [53] pathway reference for E. coli. Metabolic pathways of the co-obligate *Buchnera* lineages were compared to three *Buchnera* strains where it is the sole obligate symbiont (strains APS, G002, BAg: Table S12) to target regions that had been deleted from *Buchnera* that are in pathways involved in synthesising essential nutrients. Where genes involved in nutrient pathways were deleted in *Buchnera*, we confirmed complementary copies of the deleted genes had been retained in the *Serratia* genome. The presence of genes that are part of nutrient pathways in each *Buchnera* and *Serratia* genome was evaluated using Pathway Tools [54] and manual examination of the annotation files. See Table S6 for full results.

Additionally, in order to investigate what genes might be missing from the *A. urticata* strain of *Buchnera*, we used Orthofinder v2.2.7 [55] with default parameters to group orthologous genes between all of the *Buchnera* strains investigated (as listed in Table S12).

### Fluorescent in situ hybridisation

Whole mount FISH was performed following a protocol adapted from [56]. Aphids were fixed overnight in Carnoy’s solution (60 % ethanol, 30 % chloroform, 10 % glacial acetic acid). The aphids were then bleached in an alcoholic H_2_O_2_ solution (80 % ethanol, 14 % H_2_O, 6 % H_2_O_2_) for 3 days, changing the solution each day. The samples were then thoroughly washed in 90 % ethanol and kept at −25°C until processed. The head of the aphids was removed to facilitate the penetration of the probes, and the samples were washed 4 times (30 minutes each) in PBSTw (Phosphate Buffer Saline with 0.02 % of Tween 20), and then 3 times (5 minutes each) in hybridisation buffer (20 mM Tris-HCl at pH 8.0, 0.9 M NaCl, 0.01% sodium dodecyl sulfate and 30% formamide). The samples were then incubated overnight at room temperature in hybridization buffer supplemented with 100 mM of each fluorescent probe, one targeting *Buchnera*, one targeting *Serratia* (Table S13). Following a washing in PBSTw, the samples were mounted on slides in vectashield hardset antifade mounting media with DAPI (to highlight the host body). Mounted samples were visualised using a Leica DMRA2 epi-fluorescent microscope. Monochrome pictures were obtained using a Hamamatsu Orca camera and the Volocity 6.3.1 software, and final colour images were obtained using ImageJ. Probes were ordered from Eurogentec (Seraing, Belgium).

## QUANTIFICATION AND STATISTICAL ANALYSES

All statistical analyses were performed using R 3.6.0 [57]. Lifetime fecundity were analysed using GLMs with a quasi-Poisson distribution. Symbiont density data were analysed using GLMs with a Gamma distribution.

## DATA AND CODE AVAILABILITY

### SUPPLEMENTAL INFORMATION

Supplemental information includes one pdf document containing Figures S1-S4 and Tables S1-S5, S7-S8, S10-11 and S13 (Supplemental Information Main) and one excel document containing Tables S6, S9 and S12 (Supplemental Information Appendix).

## REFERENCES

1. McCutcheon, J.P., McDonald, B.R., and Moran, N.A. (2009). Convergent evolution of metabolic roles in bacterial co-symbionts of insects. Proc. Natl. Acad. Sci. U. S. A. 106, 15394–15399.

2. McCutcheon, J.P., and Von Dohlen, C.D. (2011). An interdependent metabolic patchwork in the nested symbiosis of mealybugs. Curr. Biol. 21, 1366–1372. Available at: http://dx.doi.org/10.1016/j.cub.2011.06.051.

3. Takiya, D.M., Tran, P.L., Dietrich, C.H., and Moran, N.A. (2006). Co-cladogenesis spanning three phyla: Leafhoppers (Insecta: Hemiptera: Cicadellidae) and their dual bacterial symbionts. Mol. Ecol. 15, 4175–4191.

4. Wu, D., Daugherty, S.C., Van Aken, S.E., Pai, G.H., Watkins, K.L., Khouri, H., Tallon, L.J., Zaborsky, J.M., Dunbar, H.E., Tran, P.L., et al. (2006). Metabolic Complementarity and Genomics of the Dual Bacterial Symbiosis of Sharpshooters. PLoS Biol. 4, e188.

5. Rao, Q., Rollat-Farnier, P.A., Zhu, D.T., Santos-Garcia, D., Silva, F.J., Moya, A., Latorre, A., Klein, C.C., Vavre, F., Sagot, M.F., et al. (2015). Genome reduction and potential metabolic complementation of the dual endosymbionts in the whitefly Bemisia tabaci. BMC Genomics 16, 1–13.

6. Douglas, A.E. (2016). How multi-partner endosymbioses function. Nat. Rev. Microbiol. 14, 731–743.

7. Pérez-Brocal, V., Gil, R., Ramos, S., Lamelas, A., Postigo, M., Michelena, J.M., Silva, F.J., Moya, A., and Latorre, A. (2006). A small microbial genome: The end of a long symbiotic relationship? Science (80-.). 314, 312–313.

8. Lamelas, A., Pérez-Brocal, V., Gómez-Valero, L., Gosalbes, M.J., Moya, A., and Latorre, A. (2008). Evolution of the secondary symbiont “Candidatus Serratia symbiotica” in aphid species of the subfamily Lachninae. Appl. Environ. Microbiol. 74, 4236–4240.

9. Manzano-Marín, A., Simon, J.C., and Latorre, A. (2016). Reinventing the wheel and making it round again: Evolutionary convergence in Buchnera-serratia symbiotic consortia between the distantly related Lachninae aphids Tuberolachnus salignus and Cinara cedri. Genome Biol. Evol. 8, 1440–1458.

10. Manzano-Marín, A., and Latorre, A. (2014). Settling down: The genome of serratia symbiotica from the aphid cinara tujafilina zooms in on the process of accommodation to a cooperative intracellular life. Genome Biol. Evol. 6, 1683–1698.

11. Henry, L.M., Maiden, M.C.J., Ferrari, J., and Godfray, H.C.J. (2015). Insect life history and the evolution of bacterial mutualism. Ecol. Lett. 18, 516–525.

12. Manzano-Marín, A., Szabó, G., Simon, J.C., Horn, M., and Latorre, A. (2017). Happens in the best of subfamilies: establishment and repeated replacements of co-obligate secondary endosymbionts within Lachninae aphids. Environ. Microbiol. 19, 393–408.

13. Meseguer, A.S., Manzano-Marín, A., Coeur d’Acier, A., Clamens, A.L., Godefroid, M., and Jousselin, E. (2017). Buchnera has changed flatmate but the repeated replacement of co-obligate symbionts is not associated with the ecological expansions of their aphid hosts. Mol. Ecol. 26, 2363–2378.

14. Nakabachi, A., and Ishikawa, H. (1999). Provision of riboflavin to the host aphid, Acyrthosiphon pisum, by endosymbiotic bacteria, Buchnera. J. Insect Physiol. 45, 1–6.

15. Douglas, A.E., and Prosser, W.A. (1992). Synthesis of the essential amino acid tryptophan in the pea aphid (Acyrthosiphon pisum) symbiosis. J. Insect Physiol. 38, 565–568.

16. Douglas, A.E. (1988). Sulphate utilization in an aphid symbiosis. Insect Biochem. 18, 599–605.

17. Febvay, G., Liadouze, I., Guillaud, J., and Bonnot, G. (1995). Analysis of energetic amino acid metabolism in Acyrthosiphon pisum: A multidimensional approach to amino acid metabolism in aphids. Arch. Insect Biochem. Physiol. 29, 45–69.

18. Liadouze, I., Febvay, G., Guillaud, J., and Bonnot, G. (1996). Metabolic fate of energetic amino acids in the aposymbiotic pea aphid Acyrthosiphon pisum (Harris) (Homoptera: Aphididae). Symbiosis 21, 115–127.

19. Brinza, L., Viñuelas, J., Cottret, L., Calevro, F., Rahbé, Y., Febvay, G., Duport, G., Colella, S., Rabatel, A., Gautier, C., et al. (2009). Systemic analysis of the symbiotic function of Buchnera aphidicola, the primary endosymbiont of the pea aphid Acyrthosiphon pisum. Comptes Rendus - Biol. 332, 1034–1049. Available at: http://dx.doi.org/10.1016/j.crvi.2009.09.007.

20. Wilson, A.C.C., Ashton, P.D., Calevro, F., Charles, H., Colella, S., Febvay, G., Jander, G., Kushlan, P.F., MacDonald, S.J., Schwartz, J.F., et al. (2010). Genomic insight into the amino acid relations of the pea aphid, Acyrthosiphon pisum, with its symbiotic bacterium Buchnera aphidicola. Insect Mol. Biol. 19, 249–258.

21. Shigenobu, S., and Wilson, A.C.C. (2011). Genomic revelations of a mutualism: The pea aphid and its obligate bacterial symbiont. Cell. Mol. Life Sci. 68, 1297–1309.

22. Akashi, H., and Gojobori, T. (2002). Metabolic efficiency and amino acid composition in the proteomes of Escherichia coli and Bacillus subtilis. Proc. Natl. Acad. Sci. U. S. A. 99, 3695–3700.

23. McCutcheon, J.P., and Moran, N.A. (2010). Functional convergence in reduced genomes of bacterial symbionts spanning 200 my of evolution. Genome Biol. Evol. 2, 708–718.

24. Douglas, A.E., and Smith, D.C. (1989). Are endosymbioses mutualistic? Trends Ecol. Evol. 4, 350–352.

25. De Mazancourt, C., Loreau, M., and Dieckmann, U. (2005). Understanding mutualism when there is adaptation to the partner. J. Ecol. 93, 305–314.

26. Kremer, N., Voronin, D., Charif, D., Mavingui, P., Mollereau, B., and Vavre, F. (2009). Wolbachia interferes with ferritin expression and iron metabolism in insects. PLoS Pathog. 5.

27. Moné, Y., Monnin, D., and Kremer, N. (2014). The oxidative environment: A mediator of interspecies communication that drives symbiosis evolution. Proc. R. Soc. B Biol. Sci. 281.

28. Bennett, G.M., and Moran, N.A. (2015). Heritable symbiosis: The advantages and perils of an evolutionary rabbit hole. Proc. Natl. Acad. Sci. U. S. A. 112, 10169–10176.

29. Sudakaran, S., Kost, C., and Kaltenpoth, M. (2017). Symbiont Acquisition and Replacement as a Source of Ecological Innovation. Trends Microbiol. 25, 375–390. Available at: http://dx.doi.org/10.1016/j.tim.2017.02.014.

30. Frank, S.A. (1997). Models of symbiosis. Am. Nat. 150, 80–99.

31. Foottit, R.G., Maw, H.E.L., Von Dohlen, C.D., and Hebert, P.D.N. (2008). Species identification of aphids (Insecta: Hemiptera: Aphididae) through DNA barcodes. Mol. Ecol. Resour. 8, 1189–1201.

32. Heie, O.E., and others (1980). The Aphidoidea (Hemiptera) of Fennoscandia and Denmark. I. General Part. The families Mindaridae, Hormaphididae, Thelaxidae, Anoeciidae, and Pemphigidae.

33. Ferrari, J., West, J.A., Via, S., and Godfray, H.C.J. (2011). Population genetic structure and secondary symbionts in host-associated populations of the pea aphid complex. Evolution (N. Y). 66, 375–390.

34. Caporaso, J.G., Lauber, C.L., Walters, W.A., Berg-Lyons, D., Huntley, J., Fierer, N., Owens, S.M., Betley, J., Fraser, L., Bauer, M., et al. (2012). Ultra-high-throughput microbial community analysis on the Illumina HiSeq and MiSeq platforms. ISME J. 6, 1621–1624.

35. Schloss, P.D., Westcott, S.L., Ryabin, T., Hall, J.R., Hartmann, M., Hollister, E.B., Lesniewski, R.A., Oakley, B.B., Parks, D.H., Robinson, C.J., et al. (2009). Introducing mothur: Open-source, platform-independent, community-supported software for describing and comparing microbial communities. Appl. Environ. Microbiol. 75, 7537–7541.

36. Quast, C., Pruesse, E., Yilmaz, P., Gerken, J., Schweer, T., Yarza, P., Peplies, J., and Glöckner, F.O. (2013). The SILVA ribosomal RNA gene database project: Improved data processing and web-based tools. Nucleic Acids Res. 41, 590–596.

37. Koga, R., Tsuchida, T., Sakurai, M., and Fukatsu, T. (2007). Selective elimination of aphid endosymbionts: Effects of antibiotic dose and host genotype, and fitness consequences. FEMS Microbiol. Ecol. 60, 229–239.

38. Lamelas, A., Gosalbes, M.J., Manzano-Marín, A., Peretó, J., Moya, A., and Latorre, A. (2011). Serratia symbiotica from the aphid Cinara cedri: A missing link from facultative to obligate insect endosymbiont. PLoS Genet. 7.

39. Madeira, F., Park, Y.M., Lee, J., Buso, N., Gur, T., Madhusoodanan, N., Basutkar, P., Tivey, A.R.N., Potter, S.C., Finn, R.D., et al. (2019). The EMBL-EBI search and sequence analysis tools APIs in 2019. Nucleic Acids Res. 47, W636–W641.

40. Guindon, S., Dufayard, J., and Lefort, V. (2010). Guindon et al. - 2010 - New Algorithms and Methods to Estimate Maximim-Likelihood Phylogenies Assessing the Performance of PhyML 3.0. Syst. Biol. 59, 307–321.

41. von Dohlen, C.D., and Moran, N.A. (1995). Molecular phylogeny of the homoptera: a paraphyletic taxon. J. Mol. Evol. 41, 211–223.

42. Rambaut, A. (2007). FigTree, a graphical viewer of phylogenetic trees.

43. Yang, Z. (2007). PAML 4: Phylogenetic analysis by maximum likelihood. Mol. Biol. Evol. 24, 1586–1591.

44. Chen, R., Favret, C., Jiang, L., Wang, Z., and Qiao, G. (2016). An aphid lineage maintains a bark-feeding niche while switching to and diversifying on conifers. Cladistics 32, 555–572.

45. Bolger, A.M., Lohse, M., and Usadel, B. (2014). Trimmomatic: A flexible trimmer for Illumina sequence data. Bioinformatics 30, 2114–2120.

46. Nurk, S., Bankevich, A., Antipov, D., Gurevich, A.A., Korobeynikov, A., Lapidus, A., Prjibelski, A.D., Pyshkin, A., Sirotkin, A., Sirotkin, Y., et al. (2013). Assembling single-cell genomes and mini-metagenomes from chimeric MDA products. J. Comput. Biol. 20, 714–737.

47. Buchfink, B., Xie, C., and Huson, D.H. (2014). Fast and sensitive protein alignment using DIAMOND. Nat. Methods 12, 59–60.

48. O’Leary, N.A., Wright, M.W., Brister, J.R., Ciufo, S., Haddad, D., McVeigh, R., Rajput, B., Robbertse, B., Smith-White, B., Ako-Adjei, D., et al. (2016). Reference sequence (RefSeq) database at NCBI: Current status, taxonomic expansion, and functional annotation. Nucleic Acids Res. 44, D733–D745.

49. Shigenobu, S., Watanabe, H., Hattori, M., Sakaki, Y., and Ishikawa, H. (2000). Genome sequence of the endocellular bacterial symbiont of aphids Buchnera sp. APS. Nature 407, 81–86.

50. Darling, A.E., Mau, B., and Perna, N.T. (2010). Progressivemauve: Multiple genome alignment with gene gain, loss and rearrangement. PLoS One 5.

51. Tanizawa, Y., Fujisawa, T., and Nakamura, Y. (2018). DFAST: A flexible prokaryotic genome annotation pipeline for faster genome publication. Bioinformatics 34, 1037–1039.

52. Riley, M., Abe, T., Arnaud, M.B., Berlyn, M.K.B., Blattner, F.R., Chaudhuri, R.R., Glasner, J.D., Horiuchi, T., Keseler, I.M., Kosuge, T., et al. (2006). Escherichia coli K-12: A cooperatively developed annotation snapshot - 2005. Nucleic Acids Res. 34, 1–9.

53. Caspi, R., Foerster, H., Fulcher, C.A., Kaipa, P., Krummenacker, M., Latendresse, M., Paley, S., Rhee, S.Y., Shearer, A.G., Tissier, C., et al. (2008). The MetaCyc Database of metabolic pathways and enzymes and the BioCyc collection of pathway/genome databases. Nucleic Acids Res. 36, 623–631.

54. Karp, P.D., Latendresse, M., Paley, S.M., Krummenacker, M., Ong, Q.D., Billington, R., Kothari, A., Weaver, D., Lee, T., Subhraveti, P., et al. (2016). Pathway tools version 19.0 update: Software for pathway/genome informatics and systems biology. Brief. Bioinform. 17, 877–890.

55. Emms, D., and Kelly, S. (2018). OrthoFinder2: fast and accurate phylogenomic orthology analysis from gene sequences. Biorxiv.

56. Koga, R., Tsuchida, T., and Fukatsu, T. (2009). Quenching autofluorescence of insect tissues for in situ detection of endosymbionts. Appl. Entomol. Zool. 44, 281–291.

57. R Core Team (2019). R: A language and environment for statistical computing.

