## Supplemental information, Main for "Parallel evolution in the integration of a co-obligate aphid symbiosis"

**Parallel evolution of a tripartite obligate symbiosis in aphids**

Table S6, S9, S12 are in the excel spreadsheet “appendix”.

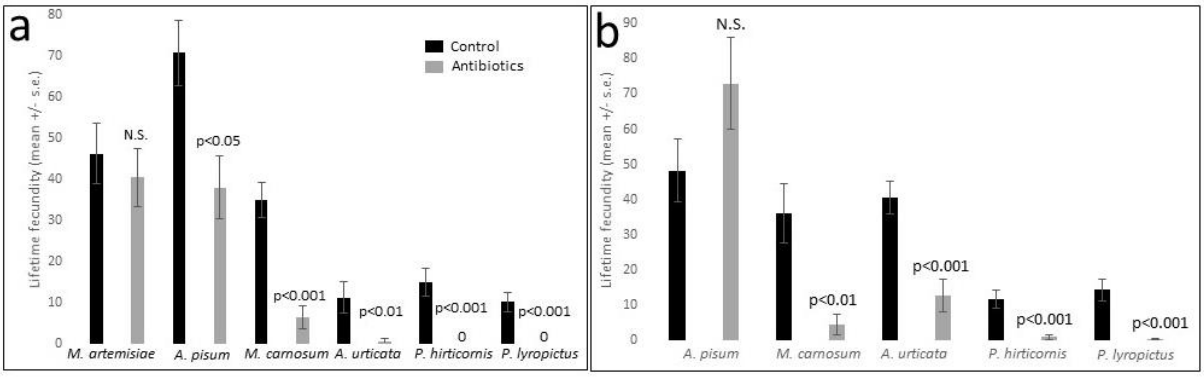


**Figure S1. Effect of antibiotic curing on aphid lifetime fecundity.** (a) Experiment 1 (United Kingdom) aphids treated for 5-days with antibiotics, and (b) Experiment 2 (Netherlands) aphids treated for 3-days with antibiotics. In both experiments, *A. pisum* carried *Serratia*. In the UK experiment, *M. artemisiae,* was included as an uninfected control.

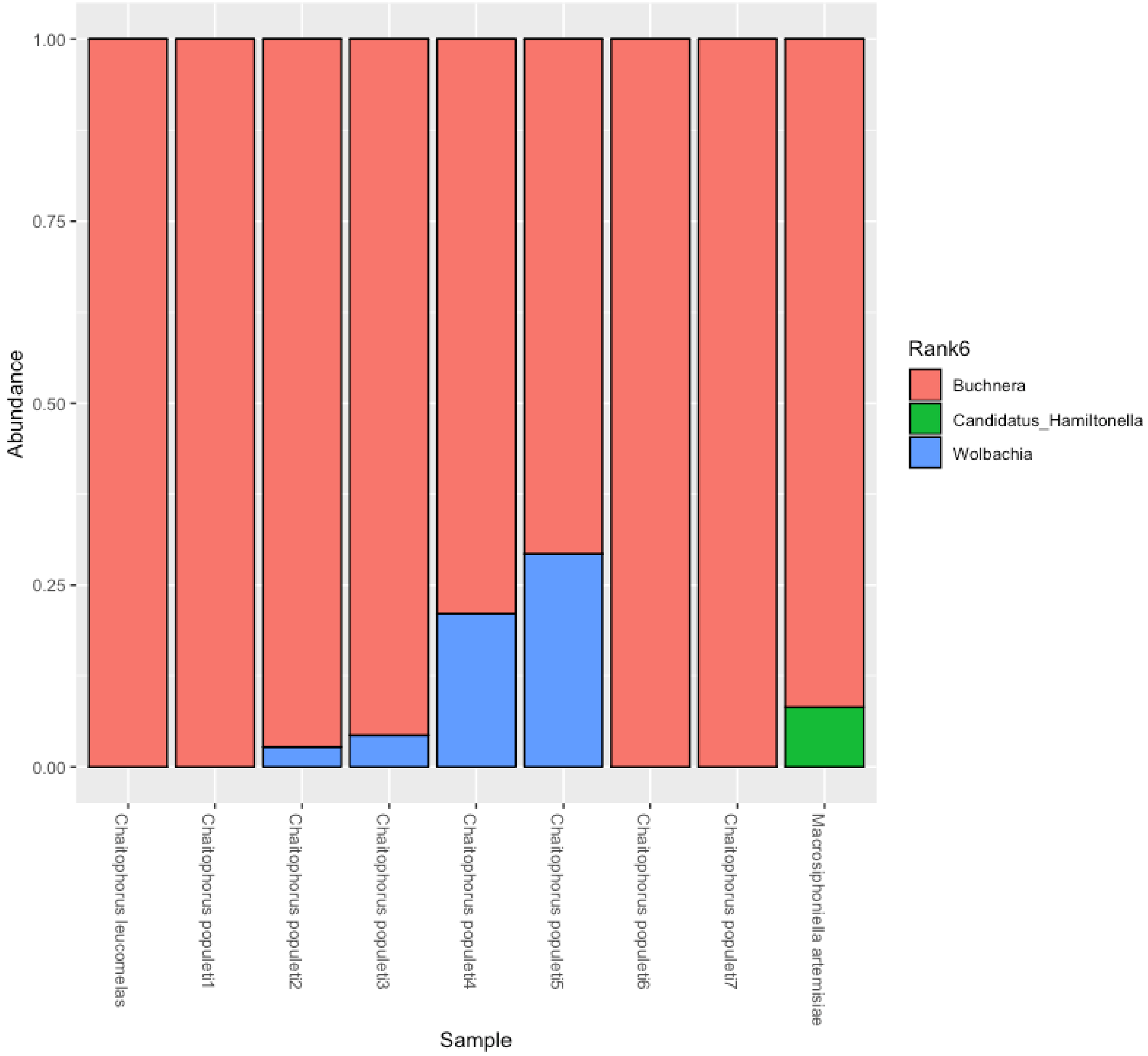


**Figure S2. Abundance of bacterial OTU's present at 1% relative abundance based on deep-coverage 16S rRNA amplicon sequencing of individual *Chaitophorus* aphids.** High prevalence of *Buchnera* and a consistent lack of *Serratia*, or any other symbionts, across individuals suggests *Buchnera* is the sole obligate symbiont in these species.

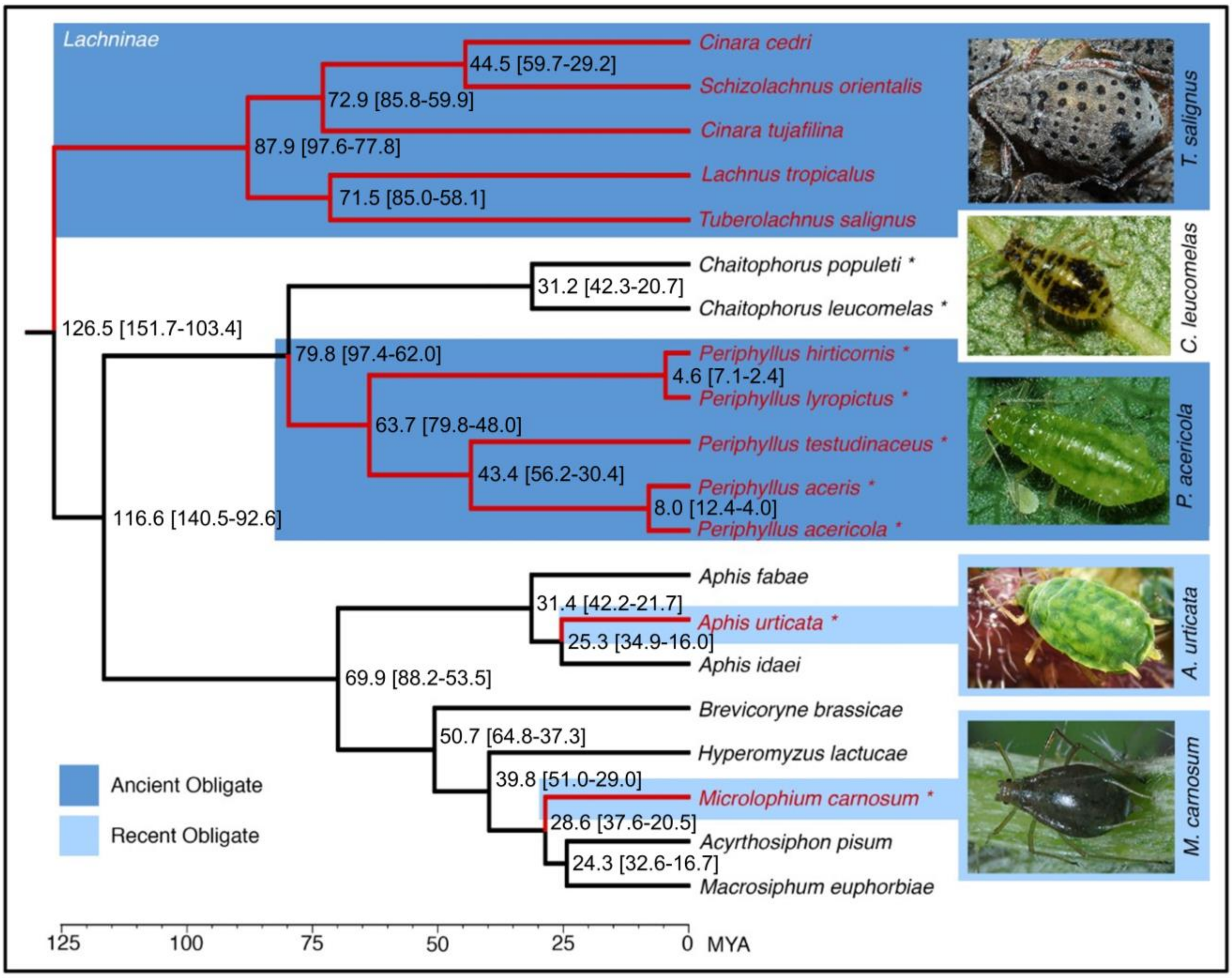


**Figure S3. Aphid phylogeny with ages based on a molecular clock.** Age estimates and 95 % confidence intervals are displayed at nodes.

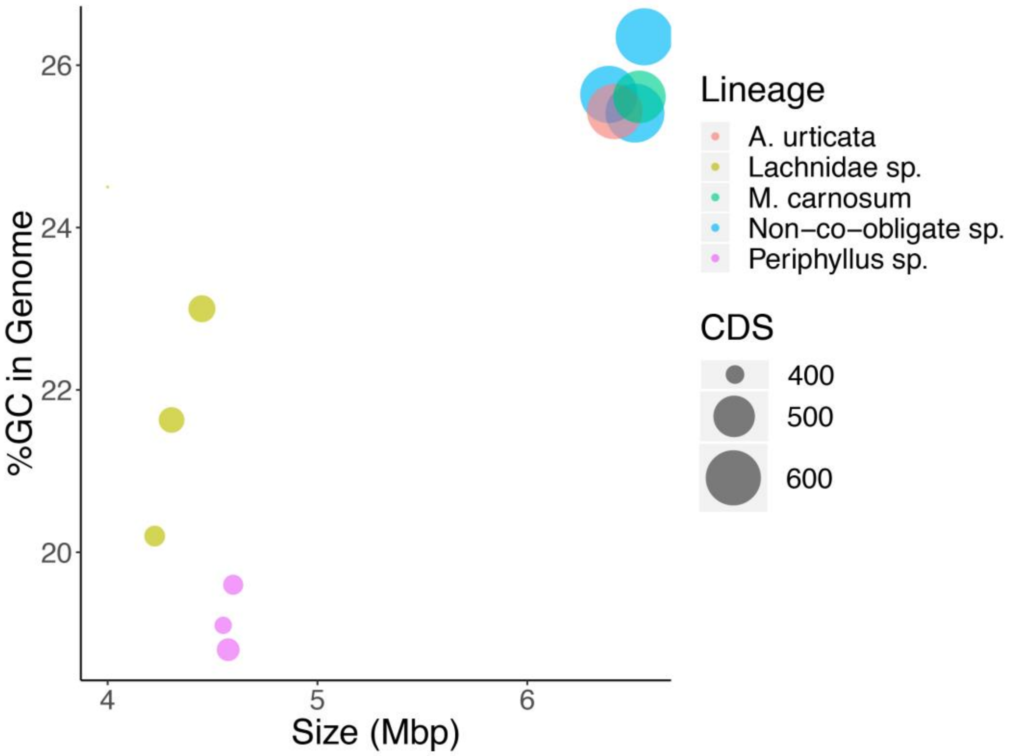


**Figure S4. Genome size, GC content and number of protein coding sequences (CDS) in *Buchnera* genomes in aphid lineages that have evolved dependency on *Serratia* compared to those that have not.** Aphid species in the Lachninae subfamily and *Periphyllus* genus have relatively small genomes, with low CDS and GC content indicative of additional genome erosion that is a product of an ancient co-obligate association with *Serratia*. Conversely, *Buchnera* from *A. urticata* and *M. carnosum* have more similar genomic features to species where *Buchnera* is the sole obligate symbionts (represented here by non-co-obligate species *A. pisum*, *M. persicae* and *A. glycines*) despite having evolved dependency on the symbiont, suggesting a more recent co-obligate relationship.

| **Aphids infected by *Serratia* / Aphids screened** | | |
|---|---|---|
| **Species** | UK aphids | Dutch aphids |
| ***A. urticata*** | 9/9 | 6/6 |
| ***M. carnosum*** | 20/20 | 7/7 |
| ***P. acericola*** | 5/5 | NA |
| ***P. aceris*** | 3/3 | NA |
| ***P. hirticornis*** | 11/11 | 20/20 |
| ***P. lyropictus*** | 9/9 | NA |
| ***P. testudinaceus*** | 70/70 | 16/16 |

**Table S1. Prevalence of *Serratia* in seven focal aphid species.** PCR screening for *Serratia* to confirm ubiquitous infections in the UK and Netherlands.

| Species | *Serratia* co-obligate | Sample size (control/treated) | Deviance | Degrees of freedom | P-value | Bonferroni corrected p-value |
|---|---|---|---|---|---|---|
| ***A. pisum*** | No | 34/36 | 35.456 | 1; 68 | 0.2734 | 1 |
| ***M. artemisiae*** | No | 16/17 | 6.3592 | 1; 31 | 0.5737 | 1 |
| ***A. urticata*** | Yes | 30/26 | 324.82 | 1; 54 | 0.0002502 | 0.0015012 |
| ***M. carnosum*** | Yes | 47/49 | 1150.5 | 1; 94 | 2.97E-10 | 1.7832E-09 |
| ***P. hirticornis*** | Yes | 58/60 | 879.77 | 1; 116 | 1.55E-10 | 9.282E-10 |
| ***P. lyropictus*** | Yes | 42/40 | 594.06 | 1; 80 | <2.2E-16 | <1.32E-15 |

**Table S2. Effect of the antibiotic treatment on lifetime fecundity of aphids (quasi-Poisson GLM**). Results from the two experiments are pooled for statistical analysis. Results of individual experiments in Fig S1.

| Experiment (origin of the aphids) | Species | Sample size (control/treated) | Deviance | Degrees of freedom | P-value | Bonferroni corrected p-value |
|---|---|---|---|---|---|---|
| **Experiment 1 (United Kingdom)** | *A. pisum* | 18/24 | 208.84 | 1; 40 | 0.006688 | 0.040128 |
| | *M. artemisiae* | 16/17 | 6.3592 | 1; 31 | 0.5737 | 1 |
| | *A. urticata* | 11/8 | 100.87 | 1; 17 | 0.001645 | 0.00987 |
| | *M. carnosum* | 35/31 | 711.57 | 1; 64 | 3.995E-07 | 0.000002397 |
| | *P. hirticornis* | 37/24 | 553.95 | 1; 59 | 1.471E-08 | 8.826E-08 |
| | *P. lyropictus* | 23/19 | 281.82 | 1; 40 | 2.888E-10 | 1.7328E-09 |
| **Experiment 2 (Netherlands)** | *A. pisum* | 16/12 | 69.641 | 1; 26 | 0.1055 | 0.5275 |
| | *A. urticata* | 19/18 | 283.22 | 1; 35 | 0.0001443 | 0.0007215 |
| | *M. carnosum* | 12/18 | 425.75 | 1; 28 | 0.0004036 | 0.002018 |
| | *P. hirticornis* | 21/36 | 304.37 | 1; 55 | 0.00002438 | 0.0001219 |
| | *P. lyropictus* | 19/21 | 332.85 | 1; 38 | 1.013E-11 | 5.065E-11 |

**Table S3. Effect of the antibiotic treatment on lifetime fecundity (quasi-Poisson GLM).** Results from each experiment are analysed separately.

| Species | Sample size (control/cured) | Deviance | Degrees of freedom | P-value | Bonferroni corrected p-value |
|---|---|---|---|---|---|
| ***M. carnosum*** | 9/11 | 28.592 | 1; 18 | 2.24E-05 | 4.48E-05 |
| ***P. hirticornis*** | 10/10 | 26.361 | 1; 18 | 3.23E-06 | 6.47E-06 |

**Table S4. Effect of the antibiotic treatment on *Serratia* density (Gamma GLM).** Results from the UK experiment.

| Species | Sample size (control/cured) | Deviance | Degrees of freedom | P-value | Bonferroni corrected p-value |
|---|---|---|---|---|---|
| ***M. carnosum*** | 9/11 | 3.0694 | 1; 18 | 0.1355 | 0.271 |
| ***P. hirticornis*** | 10/10 | 6.1222 | 1; 18 | 0.2639 | 0.5278 |

**Table S5. Effect of the antibiotic treatment on *Buchnera* density (Gamma GLM).** Results from the UK experiment.

| Gene Name | Protein Name | **Function** (as reported by UniProt for *Escherichia coli* strain K12) |
|---|---|---|
| **FliN** | Flagellar Motor Switch Protein FliN | FliN is one of three proteins (FliG, FliN, FliM) that form a switch complex that is proposed to be located at the base of the basal body. This complex interacts with the CheY and CheZ chemotaxis proteins, in addition to contacting components of the motor that determine the direction of flagellar rotation. |
| **GrpE** | Protein GrpE | Participates actively in the response to hyperosmotic and heat shock by preventing the aggregation of stress-denatured proteins, in association with DnaK and GrpE. It is the nucleotide exchange factor for DnaK and may function as a thermosensor [1]. |
| **RsxA** | Ion-translocating oxidoreductase complex subunit A | Part of a membrane-bound complex that couples electron transfer with translocation of ions across the membrane (By similarity) [2]. |
| **trpS** | Tryptophan--tRNA ligase | Catalyzes the attachment of tryptophan to tRNA(Trp). Amino acylates tRNA(Trp) with both L- and D-tryptophan, although D-tryptophan is a poor substrate [3]. |
| **YbeY** | Endoribonuclease YbeY | Single strand-specific metallo-endoribonuclease involved in late-stage 70S ribosome quality control and in maturation of the 3' terminus of the 16S rRNA [4–6]. |
| **YeeX** | UPF0265 protein YeeX | Function unknown. |

**Table S7. Identity and function of all genes that were absent from the *Buchnera* in *A. urticata* that were present in *Buchnera* from *A. pisum*, *M. persicae* and *A. glycine*, where it is the sole obligate symbiont.**

| Compound | Required | Aromatic | Cost* | Cost** | 1 | 2 | 3 | 4 |
|---|---|---|---|---|---|---|---|---|
| Tryptophan | Yes | Yes | 74.3 | ■ | ■ | ■ | | |
| Phenylalanine | Yes | Yes | 52 | ■ | ■ | ■ | ■ | |
| Histidine | Yes | Yes | 38.3 | ■ | ■ | ■ | ■ | |
| Methionine | Yes | No | 34.3 | ■ | ■ | ■ | ■ | |
| Isoleucine | Yes | No | 32.3 | ■ | ■ | ■ | ■ | |
| Lysine | Yes | No | 30.3 | ■ | ■ | ■ | ■ | ■ |
| Arginine | Yes | No | 27.3 | ■ | ■ | ■ | ■ | ■ |
| Leucine | Yes | No | 27.3 | ■ | ■ | ■ | ■ | ■ |
| Valine | Yes | No | 23.3 | ■ | ■ | ■ | ■ | |
| Threonine | Yes | No | 18.7 | ■ | ■ | ■ | ■ | ■ |

**Table S8. Metabolic Cost of Essential Amino Acid Synthesis.** Cost data is from [7], cost is as determined by number of high energy phosphate bonds contained in ATP and GTP molecules needed to form these compounds. 1 = *A. pisum* 2 = *M. carnosum* 3 = *Lachninae* subfamily species 4 = *Periphyllus* genus species.

| **Gene amplified** | **Primer names** | **Primer sequences** | **Reference** |
|---|---|---|---|
| *Buchnera dnaK* gene | BHS70F2 | 5' ATGGGTAAAATTATTGGTATTG 3' | [8] |
| | BHS70R2 | 5' ATAGCTTGACGTTTAGCAGG 3' | |
| *Serratia dnaK* gene | ApRF1 | 5'-TGGCGGGTGATGTGAAG-3' | [9] |
| | ApRR1 | 5'-CGGGATAGTGGTGTTTTTGG-3' | |
| Aphid *elongation factor 1-α* | ApEF1-alpha 107F | 5' CTGATTGTGCCGTGCTTATTG 3' | [8] |
| | ApEF1-alpha 246R | 5' TATGGTGGTTCAGTAGAGTCC 3' | |

**Table S10. Quantitative PCR primers.**

| **Gene** | **NCBI sequence identifier** | **Species** |
|---|---|---|
| Elongation factor 1 alpha (EF1a) gene | EU071358.1 | *Acyrthosiphon pisum* |
| | EF073226.1 | *Adelges cooleyi* (outgroup) |
| | EF073230.1 | *Adelges japonicus* (outgroup) |
| | EU358908.1 | *Aphis fabae* |
| | JF950580.1 | *Aphis idaei* |
| | EU358928.1 | *Brevicoryne brassicae* |
| | KX645787.1 | *Chaitophorus leucomelas* |
| | KX645797.1 | *Chaitophorus populeti* |
| | KT238076.1 | *Chaitophorus saliniger* |
| | FM174683.1 | *Cinara cedri* |
| | FM174684.1 | *Cinara tujafilina* |
| | DQ005157.1 | *Hyperomyzus lactucae* |
| | AF163879.1 | *Lachnus tropicalis* |
| | HM117788.1 | *Macrosiphum euphorbiae* |
| | KM501168.1 | *Schizolachnus orientalis* |
| | AF147812.1 | *Tuberolachnus salignus* |

| | | |
|---|---|---|
| 12S ribosomal RNA gene | AF275250.1 | *Acyrthosiphon pisum* |
| | AF275216.1 | *Adelges cooleyi* (outgroup) |
| | AF275214.1 | *Adelges japonicus* (outgroup) |
| | EU358868.1 | *Aphis fabae* |
| | EU358888.1 | *Brevicoryne brassicae* |
| | JX965987.1 | *Chaitophorus leucomelas* |
| | KX507107.1 | *Chaitophorus saliniger* |
| | KX507113.1 | *Cinara tujafilina* |
| | JX965996.1 | *Hyperomyzus lactucae* |
| | HM117803.1 | *Macrosiphum euphorbiae* |
| | AF275246.1 | *Tuberolachnus salignus* |
| 16S ribosomal RNA gene | CP034882.1 | *Buchnera aphidicola (Brevicoryne brassicae) strain Bbr* |
| | KX620606.1 | *Buchnera aphidicola (Chaitophorus leucomelas)* |
| | KX620618.1 | *Buchnera aphidicola (Chaitophorus populeti)* |
| | KX620620.1 | *Buchnera aphidicola (Chaitophorus populeti)* |
| | KX620627.1 | *Buchnera aphidicola (Chaitophorus saliniger)* |
| | CP000263.1 | *Buchnera aphidicola (Cinara cedri)* |
| | CP001817.1 | *Buchnera aphidicola (Cinara tujafilina)* |
| | CP034876.1 | *Buchnera aphidicola (Hyperomyzus lactucae) strain Hla* |
| | JX998110.1 | *Buchnera aphidicola (Lachnus tropicalis)* |
| | CP033006.1 | *Buchnera aphidicola (Macrosiphum euphorbiae) strain Meu* |
| | JX998124.1 | *Buchnera aphidicola (Schizolachnus orientalis)* |
| | LN890285.1 | *Buchnera aphidicola (Tuberolachnus salignus)* |
| | CP002301.1 | *Buchnera aphidicola str. TLW03 (Acyrthosiphon pisum)* |
| | AP010872.1 | *Candidatus Ishikawaella capsulata* |
| | CP022500.1 | *Salmonella enterica (outgroup)* |
| Cytochrome oxidase subunit 1 (COI) gene | JF883920.1 | *Acyrthosiphon pisum* |
| | KR034269.1 | *Adelges cooleyi (outgroup)* |
| | EF073073.1 | *Adelges japonicus (outgroup)* |
| | KY323028.1 | *Aphis fabae* |
| | KF638947.1 | *Aphis idaei* |
| | MH183024.1 | *Brevicoryne brassicae* |
| | KF639284.1 | *Chaitophorus leucomelas* |
| | KX680185.1 | *Chaitophorus populeti* |
| | KT237845.1 | *Chaitophorus saliniger* |
| | KU321598.1 | *Cinara cedri* |
| | JQ916729.2 | *Cinara tujafilina* |
| | KP189472.1 | *Hyperomyzus lactucae* |
| | JN032720.1 | *Lachnus tropicalis* |
| | JF883800.1 | *Macrosiphum euphorbiae* |
| | JQ916732.2 | *Schizolachnus orientalis* |
| | KT237876.1 | *Tuberolachnus salignus* |

**Table S11. Genes and sequences used in aphid phylogeny. Sequence identifiers refer to published data on NCBI.**

| Target of the probe | Sequence of the probe | 5' fluorophore | Reference |
|---|---|---|---|
| *Buchnera* 16S rRNA (*A. pisum, M. carnosum, A. urticata*) | 5'-CCTCTTTTGGGTAGATCC-3' | Alexa Fluor 488 | [10] |
| *Serratia* 16S rRNA | 5'-CCCGACTTTATCGCTGGC-3' | Cy3 | |
| *Buchnera* 16S rRNA (*Periphyllus spp.*) | 5'-CCTTTTTTGGGCAGATTC-3' | Alexa Fluor 488 | N/A |

**Table S13. Fluorescent *in situ* probes.**

### SUPPLEMENTARY REFERENCES


1. Wu, B., Wawrzynow, A., Zylicz, M., and Georgopoulos, C. (1996). Structure-function analysis of the Escherichia coli GrpE heat shock protein. EMBO J. *15*, 4806–4816.
2. Koo, M.S., Lee, J.H., Rah, S.Y., Yeo, W.S., Lee, J.W., Lee, K.L., Koh, Y.S., Kang, S.O., and Roe, J.H. (2003). A reducing system of the superoxide sensor SoxR in Escherichia coli. EMBO J. *22*, 2614–2622.
3. Soutourina, J., Plateau, P., and Blanquet, S. (2000). Metabolism of D-aminoacyl-tRNAs in Escherichia coli and Saccharomyces cerevisiae cells. J. Biol. Chem. *275*, 32535–32542.
4. Rasouly, A., Davidovich, C., and Ron, E.Z. (2010). The heat shock protein YbeY is required for optimal activity of the 30S ribosomal subunit. J. Bacteriol. *192*, 4592–4596.
5. Davies, B.W., Köhrer, C., Jacob, A.I., Simmons, L.A., Zhu, J., Aleman, L.M., RajBhandary, U.L., and Walker, G.C. (2010). Role of escherichia coli YbeY, a highly conserved protein, in rRNA processing. Mol. Microbiol. *78*, 506–518.
6. Jacob, A.I., Köhrer, C., Davies, B.W., RajBhandary, U.L., and Walker, G.C. (2013). Conserved Bacterial RNase YbeY Plays Key Roles in 70S Ribosome Quality Control and 16S rRNA Maturation. Mol. Cell *49*, 427–438.
7. Akashi, H., and Gojobori, T. (2002). Metabolic efficiency and amino acid composition in the proteomes of Escherichia coli and Bacillus subtilis. Proc. Natl. Acad. Sci. U. S. A. *99*, 3695–3700.
8. Dunbar, H.E., Wilson, A.C.C., Ferguson, N.R., and Moran, N.A. (2007). Aphid thermal tolerance is governed by a point mutation in bacterial symbionts. PLoS Biol. *5*, 1006–1015.
9. Burke, G., Fiehn, O., and Moran, N. (2010). Effects of facultative symbionts and heat stress on the metabolome of pea aphids. ISME J. *4*, 242–252.
10. Koga, R., Tsuchida, T., and Fukatsu, T. (2003). Changing partners in an obligate symbiosis: A facultative endosymbiont can compensate for loss of the essential endosymbiont Buchnera in an aphid. Proc. R. Soc. B Biol. Sci. *270*, 2543–2550.
